## Supplemental File 1 for "Genetic and pharmacological evidence for kinetic competition between alternative poly(A) sites in yeast"

**Supplemental Tables**

**Table S1 – Yeast Strains**

| **Strain** | **Genotype** | **Relevant mutation if known** | **Reference** |
| --- | --- | --- | --- |
| W303 | *MAT***a** *leu2-3,112 his3-11,15 trp1-1 can1-100 ade2-1 ura3-1* | Wild type | Thomas and Rothstein (1989) |
| BY4741 | *MAT***a** *leu2∆0 his3∆1 ura3∆0 met15∆0* | Wild type | Baker Brachmann et al. (1998) |
| *rna14-1* | *MAT***a** *leu2-3,112 his3-11,15 trp1-1 ade2-1 ura3-1 rna14-1* | T nucleotide insertion at 1986 causing a premature stop codon (replacement of last 16 amino acids with FK) | Bloch et al. (1978)(named *cor1-1*); Minvielle-Sebastia et al. (1994); Rouillard et al. (2000) |
| *rna15-1* | *MAT***a** *leu2-3,112 his3-11,15 trp1-1 ade2-1 ura3-1 rna15-1* | L214P | Bloch et al. (1978)(named *cor2-1*);  Minvielle-Sebastia et al. (1994); Qu et al. (2007) |
| *pcf11-2* | *MAT***a** *leu2-3,112 his3-11,15 trp1∆ can1-100 ade2-1 ura3-1 pcf11-2* | E232G, D280G, C424R, S538G, F562S, S579P | Amrani et al. (1997); Sadowski et al. (2003) |
| *pcf11-13* | *MAT***a** *leu2-3,112 his3-11,15 trp1∆ can1-100 ade2-1 ura3-1 pcf11-13* | D68A, S69A, I70A (CID region) | Sadowski et al. (2003) |
| *clp1-pm* | *MAT***a** *leu2-3,112 his3-11,15 trp1-1 can1-100 ade2-1 ura3-1 clp1-pm* | K136A, T137A (P-loop motif) | Ramirez et al. (2008) |
| *nab4-1* | *MAT***a** *ura3-52 hrp1-1* | N167D, F179Y, P194H, Q265L | Minvielle-Sebastia et al. (1998) |
| *nab4-1* | *MAT***a** *ura3-52 hrp1-7* | S64R, K92M, T125A, I163T | Minvielle-Sebastia et al. (1998) |
| *yhh1-3* | *MAT***a** *leu2-3,112 his3-11,15 trp1-1 can1-100 ade2-1 ura3-1 yhh1-3* | L302P, N716S, N762S, Y766C, K933R, D1070E, N1136D | Dichtl et al. (2002b) |
| *ysh1-13* | *MAT***a** *leu2-3,112 his3-11,15 trp1∆ ade2-1 ura3-1 TRP1::ysh1 [ysh1-3-HIS3-CEN]* | V235A, N685H, D695V, E723V, R763G | Garas et al. (2008) |
| *yth1-1* | *MAT***a** *leu2-3,112 his3-11,15 trp1∆ ade2-1 ura3-1 yth1::TRP1 [CEN4-ADE2-yth1-1]* | frameshift at 154 in causing a premature stop codon (deletion of last 55 amino acids) | Barabino et al. (1997) |
| *yth1-4* | *MAT***a** *leu2-3,112 his3-11,15 trp1∆ ade2-1 ura3-1 yth1::TRP1 [CEN4-ADE2 -yth1-4]* | W70A | Barabino et al. (2000) |
| *fip1-1* | *MAT***a** *leu2-3,112 his3-11,15 trp1-1 ura3-1 fip1-1* | L99F, Q216X (deletion of last 111 amino acids) | Preker et al. (1995); Preker et al. (1997) |
| *pap1-1* | *MAT***a** *ade1 ade2 lys2 gal1? ura3-52 pap1-1* |  | Patel and Butler (1992) |
| *pta1-1* | *MAT***a** *leu2-3,112 his3-11,15 trp1-1 ade2-1 ura3-1 pta1-1* | Premature stop codon | O'Connor and Peebles (1992); Preker et al. (1997) |
| *pfs2-1* | *MAT***a** *leu2-3,112 his3-11,15 trp1∆ ade2-1 ura3-1 pfs2::TRP1 [pfs2-1-LEU2-CEN]* |  | Ohnacker et al. (2000) |
| *mpe1-1* | *MAT***α** *leu2-3,112 his3-11,15 trp1-1 ade2-1 ura3-1 mpe1-1* | F9S, Q268K, K337F, K354X (deletion of last 87 amino acids) | Vo et al. (2001) |
| *ref2-2* | *MAT***a** *leu2-3,112 his3-11,15 trp1-1 can1-100 ade2-1 ura3-1 ref2-2* |  | Dheur et al. (2003) |
| *pti1-2* | *MAT***a** *leu2-3,112 his3-11,15 trp1-1 can1-100 ade2-1 ura3-1 pti1-2* |  | Dheur et al. (2003) |
| *ssu72-2* | *MAT***a** *leu2-3,112 his3-11,15 trp1-1 ade2-1 ura3-1 ssu72-2* | R129A | Dichtl et al. (2002a); Pappas and Hampsey (2000) |
| *glc7-5* | *MAT***a** *leu2-3,112 his3-11,15 can1-100 ade2-1 ura3-1 glc7::LEU2 trp1:: glc7-5* | F226L | Peggie et al. (2002) |
| *swd2-2* | *MAT***a** *leu2∆0 his3∆1 ura3∆0 met15∆0 swd2-2* | F14P, C27G, F41I, K185E, S253L, C257R | Dichtl et al. (2004) |
| *∆syc1* | *MAT***a** *leu2∆0 his3∆1 ura3∆0 met15∆0 syc1∆* | Whole gene deletion | Zhelkovsky et al. (2006) |
| *nrd1-5* | *MAT***a** *ura3-52 trp1-1 ade2-1 leu2-3,112 his3-11,15 lys2-∆2 can1-100 met2-∆1 nrd1-5* | G368V (RNA recognition motif) | Steinmetz and Brow (1996) |
| *nrd1-∆CID* | *MAT***a** *ura3-52 trp1-1 ade2-1 leu2-3,112 his3-11,15 lys2-∆2 can1-100 met2-∆1 nrd1-∆CID* | deletion of amino acids 39-169 (CTD region) | Steinmetz and Brow (1998) |
| *nab3-11* | *MAT***a** *leu2-3,112 his3-11,15 trp1-1 can1-100 ade2-1 ura3-1 nab3-11* | F371L, P374L (RNA recognition motif) | Conrad et al. (2000) |
| ***sen1-1 **** | *MAT***a** *leu2-3,112 his3-11,15 trp1-1 ade2-1 ura3-1 sen1-1* | G1747D (helicase domain) | Mischo et al. (2011); Winey and Culbertson (1988) |
| *rbp1-1* | *MAT***α** *ura3-52 leu2-3,112 rpb1-1* |  | Nonet et al. (1987) |

**Table S2 – Oligonucleotide Primers**

All primers have a concentration of 100µM unless stated

| Primer | Sequence (5’ – 3’) |
| --- | --- |
| PAT-seq biotin end-extend | Biotin-CAGACGTGTGCTCTTCCGATCTTTTTTTTTTTTTTTTTT |
| PAT-seq splint A (200µM) | CCCTACACGACGCTCTTCCG(rA)(rT)(rC)(rT) |
| PAT-seq Splint B (200µM) | NNNNAGATCGGAAGAGCGTCGTGTAGGG |
| Illumina universal Rd1 forward (50µM for mPAT) | AATGATACGGCGACCACCGAGATCTACACTCTTTCCCTACACGACGCT  CTTCCG |
| PAT anchor reverse | GCGAGCTCCGCGGCCGCGTTTTTTTTTTTT |
| TVN-PAT anchor reverse | GCGAGCTCCGCGGCCGCGTTTTTTTTTTTTVN |
| OM14 PAT forward | GGGTCTTTTGACGCTGGAC |
| mPAT reverse | CAGACGTGTGCTCTTCCGATCTTTTTTTTTTTT |
| mPAT-TVN reverse | CAGACGTGTGCTCTTCCGATCTTTTTTTTTTTTVN |
| AAD16 mPAT | CCTACACGACGCTCTTCCGATCTTGCGTTTGTGTAAGAAATATGC |
| ADE2 mPAT | CCTACACGACGCTCTTCCGATCTAGAAACTGTCGGTTACGAAGC |
| APQ12 mPAT | CCTACACGACGCTCTTCCGATCTGAAACGCCTCTGCTTACTCGG |
| ARG8 mPAT | CCTACACGACGCTCTTCCGATCTGGCTATTGAAGCGGTTTACG |
| ARP5 mPAT | CCTACACGACGCTCTTCCGATCTGAGACAGCAAACTGAAACGC |
| CHD1 mPAT | CCTACACGACGCTCTTCCGATCTGCTGATGGCAATGTACGAC |
| COX17 mPAT | CCTACACGACGCTCTTCCGATCTCTGACAGTCTGCCGACAACCA |
| CRN1 mPAT | CCTACACGACGCTCTTCCGATCTCGGCGGCGATAATAATGC |
| CST6 mPAT | CCTACACGACGCTCTTCCGATCTCTCGAGCTGCATCCTTTCTT |
| DBF2 mPAT | CCTACACGACGCTCTTCCGATCTTCAACTAGCACCTATGAACGC |
| ECM16 mPAT | CCTACACGACGCTCTTCCGATCTGCTTCCAGACCATCACAGG |
| ECM25 mPAT | CCTACACGACGCTCTTCCGATCTGCATATACGACAACAAAATACCC |
| END3 mPAT | CCTACACGACGCTCTTCCGATCTGCAGAAATCAATTGACACCGA |
| ENT1 mPAT | CCTACACGACGCTCTTCCGATCTGTGATTCTGTCATTCCAGTCCG |
| ERG8 mPAT | CCTACACGACGCTCTTCCGATCTGGTAGATAATAGTGGTCCATGTGA |
| GFD1 mPAT | CCTACACGACGCTCTTCCGATCTCACATGGACACTTTTAAGCACG |
| HSP26 mPAT | CCTACACGACGCTCTTCCGATCTGGTTTCTTCTCAAGAATCGTG |
| IMP2 mPAT | CCTACACGACGCTCTTCCGATCTGAGCCATTTTAGAATGAAAATCAGC |
| LOS1 mPAT | CCTACACGACGCTCTTCCGATCTGCAAGGTCAATAGCTTTCAGG |
| MRPL19 mPAT | CCTACACGACGCTCTTCCGATCTCGAGAAATTTTCAACAGACCTTCC |
| MRPL22 mPAT | CCTACACGACGCTCTTCCGATCTGCTGAGAAAGATGAACTGCTACTC |
| MSA1 mPAT | CCTACACGACGCTCTTCCGATCTGCATGTGAATGGAGTTGACCTTC |
| NOP16 mPAT | CCTACACGACGCTCTTCCGATCTGGCAACTACCAATTGATTACCA |
| NOT3 mPAT | CCTACACGACGCTCTTCCGATCTCAGTGCTAATGGCAGTATAATTTG |
| NUP159 mPAT | CCTACACGACGCTCTTCCGATCTGCTATATGTACGTTGTTAGTGCCG |
| OM14 mPAT | CCTACACGACGCTCTTCCGATCTGGTCTTTTGACGCTGGACGG |
| OM45 mPAT | CCTACACGACGCTCTTCCGATCTCTGGAGCTCGAAAAAGGAC |
| PDE2 mPAT | CCTACACGACGCTCTTCCGATCTTTCCTTTTGTGAAGTATTTGTGC |
| PNT1 mPAT | CCTACACGACGCTCTTCCGATCTCATGACATTATCTATGCTGTACATATTG |
| PRY2 mPAT | CCTACACGACGCTCTTCCGATCTGGATTCTTCTTTTCTAGGGTACGC |
| RCL1 mPAT | CCTACACGACGCTCTTCCGATCTGGTGTAACTTCACGGACAACT |
| RER2 mPAT | CCTACACGACGCTCTTCCGATCTGAGCAAGATAAATGAGTTCGC |
| RHO1 mPAT | CCTACACGACGCTCTTCCGATCTCAATCCCATTCCTTTTCTCA |
| RPF1 mPAT | CCTACACGACGCTCTTCCGATCTCCGTAAGAACCGTGGTCG |
| RSC4 mPAT | CCTACACGACGCTCTTCCGATCTGTCTTCCCATCATATGCATGT |
| SLF1 mPAT | CCTACACGACGCTCTTCCGATCTGGTGAAATTAGCAGGCAGTTTG |
| SNF2 mPAT | CCTACACGACGCTCTTCCGATCTGCATGACAGAAGCGAGTGTATAG |
| SRP68 mPAT | CCTACACGACGCTCTTCCGATCTGGTTTCTTGGGCCTATTTGG |
| TIM54 mPAT | CCTACACGACGCTCTTCCGATCTCCAAGGAAGAGCCAGAATCAG |
| TOM70 mPAT | CCTACACGACGCTCTTCCGATCTGCAGCAATGACATTGACATCTCAC |
| TRP2 mPAT | CCTACACGACGCTCTTCCGATCTAATGATGTATAGCAGGATCCTGA |
| UBC9 mPAT | CCTACACGACGCTCTTCCGATCTGAATCCATCTTTCCCATTCTTCC |
| VTS1 mPAT | CCTACACGACGCTCTTCCGATCTGCATATACCAACACAGGGAACA |
| YET3 mPAT | CCTACACGACGCTCTTCCGATCTGTCGATGTGCAAAAGCCTACA |
| YRA1 mPAT | CCTACACGACGCTCTTCCGATCTACCGCCACTAGGTGACGC |
| YSC84 mPAT | CCTACACGACGCTCTTCCGATCTGGATGGGTTCCTTATTCAGC |
| MID2 mPAT | CCTACACGACGCTCTTCCGATCTCAAGGTAACGAATTATCACCACG |
